# Ambient Light Impacts Innate Behaviors of New-World and Old-World Mice

**DOI:** 10.1101/2025.05.14.653927

**Authors:** Katja Reinhard, Rani Vastenavondt, Benjamin Crevits, Sybren De Boever, Po-Yu Liao, Lorenza Tortella, Karl Farrow

## Abstract

Animals encounter predators and prey under diverse lighting conditions that signal different risks and opportunities^1–4^, yet how ambient illumination shapes innate approach and avoidance behaviors remains poorly understood. Here we have systematically tested the visually guided behaviors of New-World (North American *Peromyscus*) and Old-World (Eurasian *Mus*) mice under conditions mimicking bright daylight or dim moonlit environments. We identified common and species-specific adaptations to the different lighting conditions. Across species, dim light enhanced the speed and vigor of escape responses to predator-like looming stimuli. However, species diverged in their reactions to non-threatening stimuli, with *Mus musculus* and *Peromyscus polionotus* increasing aversive behaviors under dim light, while *Peromyscus maniculatus* showed stronger avoidance under bright conditions. Finally, although ambient light levels had a common effect on exploratory behavior, these changes were not predictive of subsequent visually evoked behaviors. Our findings reveal that ambient lighting profoundly and differentially shapes innate behavioral strategies across species, and demonstrate that these context-specific survival responses are inherited rather than learned.

## INTRODUCTION

In many vertebrates, including small rodents, avoiding threat and hunting prey is guided by vision^5– 7^. In nature, rodents perform these essential behaviors under varying light conditions, which are commonly associated with different levels of risk of being attacked, or opportunities to find food. On the other hand, the behavior and success of predators also changes with lighting conditions. For instance, owls seem to be more efficient at capturing mice during a full-moon in comparison to darker nights^8–10^. Most small rodents, including species of *Mus* and *Peromyscus* are primarily active during the hours of darkness at night^11^, but also forage in brighter conditions during the hours of dawn and dusk. Independent of the time of day, mice experience a variety of lighting conditions that are dependent on where they are foraging (open exposed locations vs in the shadows), the weather conditions, and seasonal variations. At the same time, ambient light levels alter the foraging behavior with each species showing its own preference for increasing or decreasing foraging behavior in the presence of natural moon or artificial light^12–14^. In laboratory settings, innate aversive and orienting behavioral experiments are commonly performed using computer monitors that result in environments with photopic light levels similar to those experienced in indoor spaces or in the shade on a bright sunny day. Hence, our knowledge about visually-guided innate behaviors is limited to light conditions that exist at dawn, dusk, a cloudy day or when moving through dense vegetation. The effect of ambient light on behaviors such as escape or pursuit, and whether context-specific behaviors are learned or inherited, however, remains unclear.

## RESULTS

### Old-World and New-World mice display a similar set of defensive behaviors

We set out to characterize and compare the visually triggered innate defensive behaviors of three rodent species under different lighting conditions. The rodents include two species of New-World mice, genus *Peromyscus* (*Peromyscus maniculatus bairdii*; N = 39 mice, subsequently referred to as *P. maniculatus;* and *Peromyscus polionotus subgriseus;* N = 41 mice, referred to as *P. polionotus*), and one Old-World rodent species (*Mus musculus*, C57Bl/6J; N = 26 mice, subsequently called *M. musculus*) (Suppl. Figure 1A). To assess their innate reaction to overhead visual threats, mice were placed in a box with two screens, one on the top and one along the long side, with a shelter placed at one end (Figure 1A). A session began by placing one mouse in the arena and allowing it to acclimatize for a period of 10 minutes, after which one out of four visual “threat” stimuli was randomly presented on the screen above the arena each time the mouse entered the “threat zone”. Each session included no more than one presentation of each stimulus, which included either: 10 consecutive repetitions of a black looming disk (black looming), 5 repetitions of a white looming disk (white looming), 5 repetitions of a dimming disk (dimming) or a single presentation of a sweeping small disk (sweeping; see Methods, Figure 1B). Mice were filmed from below through the red plexiglass floor and DeepLabCut was used in combination with manual curation to track the mice and characterize their behavioral responses^15,16^.

**Fig. 1:**
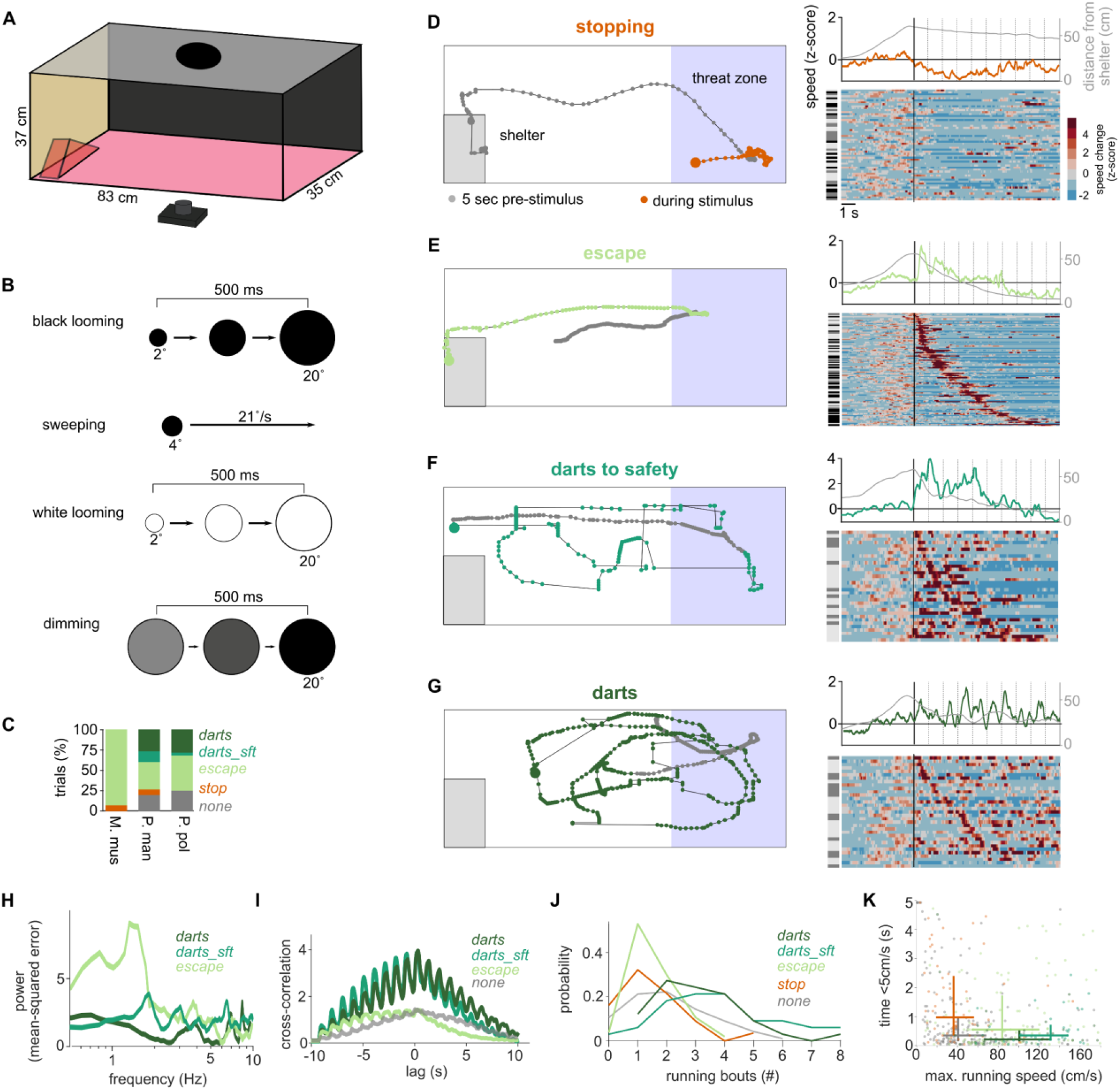
Innate aversive behaviors in mice. (**A**) Experimental setup for threat stimuli. (**B**) Visual threat stimuli. (**C**) Percentage of stopping, escape, darting to safety and other darts in response to black looming under bright light conditions. *M. musculus* (N = 14 mice), *P. maniculatus* (N = 14 mice), *P. polionotus* (N = 25 mice). (**D**) Example trajectory of a mouse reacting with stopping to a visual threat (left; see Suppl. Video 1). Right: All stopping response to any of the four overhead stimuli at all light levels and of all three tested species. Each row corresponds to a reaction by one mouse to one stimulus. Rows are sorted by onset of stopping. Gray bars on the left indicates the species for each reaction (*M. musculus*: black, *P. maniculatus*: dark gray, *P. polionotus*: light gray). Speed is given as z-score calculated from mean and standard deviation of the pre-stimulus movement speed. Above the heatmap: Mean speed change across all rows (orange) and mean distance from the shelter (gray). (**E**) Example trajectory of a mouse reacting with escape (Suppl. Video 2) and all escape trials sorted by escape onset. (**F**) Example trajectory of a mouse reacting with darts to safety (Suppl. Video 3) and all trials sorted by escape onset. (**G**) Example trajectory of a mouse reacting with darts not ending in safety (Suppl. Video 4) and all trials sorted by escape onset. (**H**) Power spectrum of the mean Fourier Transform of the running speed during darts (dark green; n = 27), darts to safety (green; n = 29) and escape (light green; n = 88) in response to any of the repetitive stimuli. (**I**) Cross correlation between running speed and visual stimulus for different behavior types and for all repetitive stimuli. Same trials as in I were included; no-behavior trials (gray; n = 106). (**J**) Numbers of running bouts for darts, darts to safety, escape, stopping (orange) and no behavior. (**K**) Maximal running speed and time spent without movement (<5cm/s) for trials classified as no behavior (gray), stopping (orange), escaping (light green) and darts to safety (green) and other darts (dark green). Lines indicate 25% to 75% quantiles and cross at the median.

The behavior of the mice was classified into one of four categories: stopping/freezing, running/escape, darting to safety and darts that did not end in safety (see Methods for definitions). Under standard experimental conditions (native monitor brightness), the typical avoidance behaviors (freezing and escape) were exhibited by each species (Figure 1C-E; Suppl. Figure 1C; Suppl. Video 1-2). In the two species of *Peromyscus*, we observed an additional darting behavior, which consisted of a series of sprints across the arena (Figure 1F-G), similar to behavior that has been described previously in some mouse strains^17^. We separated darting trials into those that ended in safety (darts_sft’; ending at the short ends of the setup, Figure 1F; Suppl. Figure 1B; Suppl. Video 3) and those that did not (Figure 1G; Suppl. Video 4). In *Mus* literature, escape behaviors are often defined as crossing of a speed threshold and ending in a shelter, but do not necessarily describe the precise trajectory. Such behaviors might be included both in our escape group (all escaping animals visited the safety zone) and in our ‘darts to safety’, although we did not observe any such behavior in *Mus* in our data set (Suppl. Figure 1C). Instead, darts of either type were most commonly observed in *Peromyscus* responses to the black looming stimulus (Figure 1C and Suppl. Figure 1C; 6 out of 14 trials for *P. maniculatus* and 9 out of 25 trials for *P. polionotus*). The fast, non-directed sprints that characterize darting behavior tended to be aligned to the onset of each loom with pauses or intermediate sprints during the time between looms. Because of this stimulus-locked behavior, oscillations can be observed in the mean speed for each bout of darting behavior (Figure 1F and 1G top). This is also evident both in the peak around 1 Hz in the Fourier analysis of darts to safety trials, which matches the frequency of the looming and dimming stimuli (Figure 1H), and the strong cross-correlation between the mice’ running speed and visual stimulation for lags of up to 10s (the duration of black looming stimuli) (Figure 1I). This stimulus locked behavior was not evident in the escape and no-behavior trials. The differences among the behavioral types were confirmed by automatic extraction of running bouts in each behavior category (Figure 1J) and by comparing the maximum running speed with the duration of time spent sitting still (Figure 1K and Suppl. Fig. 1D). Taken together, parameterization of the behavior fits well our manual classification and matches the expected running-like responses for repeated full-contrast black looming stimuli at bright ambient light levels as previously reported both in *Mus*^5^ and *Peromyscus*^18^.

### Earlier and more vigorous response to looming under dim light conditions

To investigate the impact of ambient light on defensive behaviors, we exposed mice to the same set of visual stimuli at two light levels: 128 lux, bright light conditions, and 0.2 lux, dim light conditions (see Methods; Figure 1 includes data from both conditions). The bright light condition corresponds to photopic environments experienced during a cloudy day, while the dim light condition reflects light levels experienced during a moonlit night. The order of exposure to the two lighting conditions was randomized such that approximately half of the mice were first exposed to dim conditions, and the other half to bright conditions, with a minimum of 5 days between the two sessions.

The most striking behavioral differences observed between the two lighting conditions were in response to the black looming stimulus. In all three species, we observed earlier and more vigorous responses under dim light conditions (*M. musculus:* Suppl. Video 2 vs. Suppl. Video 5; *P. maniculatus:* Suppl. Video 6 vs. Suppl. Video 7; *P. polionotus:* Suppl. Video 8 vs Suppl. Video 3). In each species, this was evident in the higher running speeds and the earlier onset of running bouts exhibited in the dim light condition (Figure 2A-B). Comparable trends were observed in *M. musculus* for experiments performed at different institutes and in a different mouse line (Suppl. Fig. 2A-B). The increased vigor of running behaviors at dimmer light levels was evident in the higher maximal running speed (Figure 2C), and the more frequent occurrence of dart behaviors in each species of *Peromyscus* (*P. polionotus*: dim 68% vs bright 39%; *P. maniculatus*: 60% vs 40%; Figure 2D), which is also reflected in the stronger cross-correlation of their behavior at dim light levels with the black looming stimulus (*P. maniculatus* r = 0.26/0.18, (dim/bright), *P. polionotus* r = 0.13/0.09; Figure 2A right). In all three species, running-like behaviors (escaping, darts to safety and other darts) started earlier under dim light conditions (Figure 2E).

**Fig. 2:**
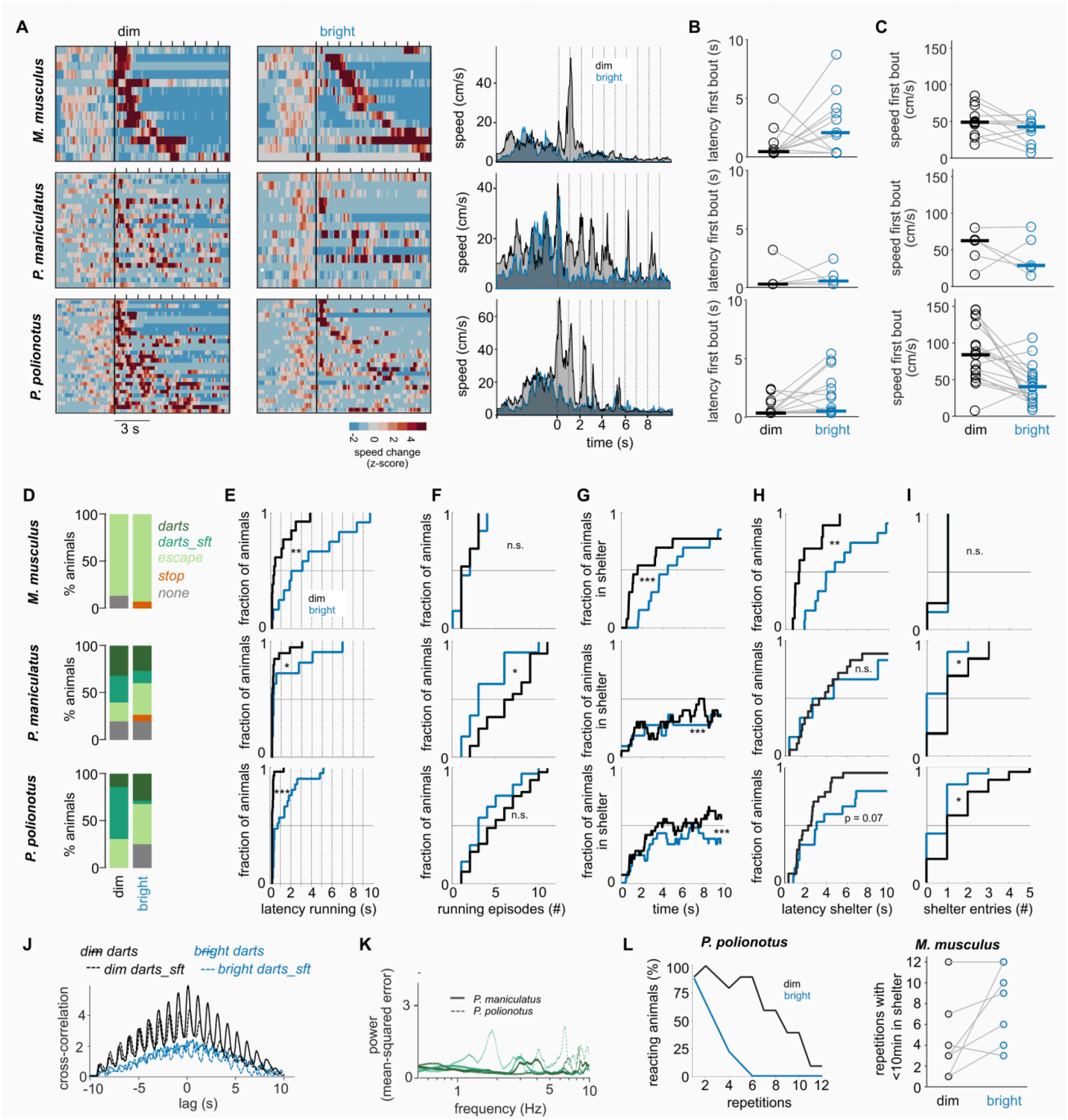
Innate reactions to black looming are more vigorous under dim compared to bright conditions. (**A**) Speed changes in response to a black looming stimulus. Each row represents the response of one mouse during the visual stimulus. Running speed is normalized to the mean and standard-deviation of the speed in the 5 seconds before stimulus onset. Ticks on top of each heatmap indicate stimulus onsets. Mean absolute running speed for all mice is shown on the right for dim and bright conditions. *Mus*: N = 15/14 (dim/bright). *P. maniculatus*: N = 25/15. *P. polionotus*: N = 29/28. (**B**) Latency of first running bout (>30cm/s) for the subset of mice that were tested and displayed running bouts at both light levels. Horizonal lines indicate medians. Mm: N = 9, Pm: N = 5, Pp: N = 19. (**C**) Maximum speed during first running bout for the same mice as in B. (**D**) Behavior types under dim and bright light. (**E-I**) Comparison of various parameters for running-like trials only (escape, darts to safety, darts). (**E**) Distribution of latency to running onset. (**F**) Number of running episodes. (**G**) Fraction of mice in the shelter during looming stimuli. (**H**) Latency to fist shelter entry. (**I**) Number of shelter entries. (**J**) Cross-correlation of darts in response to the black looming stimulus for dim and bright light conditions separately. (**K**) Power spectrum of Fourier transform for dart trials of *P. maniculatus* (solid; N = 7 for darts to safety, green, and N = 8 for other darts, dark green) and *P. polionotus* (dashed, N = 18 / N = 5) at dim light conditions. (**L**) Left: Percentage of *P. polionotus* tested and reacting to 12 repetitions of the looming stimulus (percentage indicates fraction of animals that did not stay for >10 min under the shelter during the previous repetition and showed a reaction to the looming stimulus; 100% corresponds to N = 10 mice). Right: Number of looming repetitions that could be tested in each *M. musculus* (N = 7) without inducing prolonged shelter visits of >10 min. * p < 0.05, ** p < 0.01, *** p < 0.005 Permutation Test.

Although response vigor increased in each species under dim light conditions, we identified several differences between the species. For instance, independently of the light levels, most *M. musculus* individuals showed fewer running episodes to repeated looming than the two species of *Peromyscus* and ran more directly to the shelter (Figure 2F-H; 12 out of 13 escaping *M. musculus* enter the shelter at dim light levels; 12 out of 13 at bright light) from where they did not exit for the remaining duration of the stimulus (Figure 2I). The *Peromyscus* mice visited the shelter as well, but did leave again during the same stimulus presentation (Figure 2G and 2I). Similar numbers of *P. polionotus* visited the shelter at both light levels (dim: 24 out of 29 escaping/darting mice; bright: 15 out of 21), but did so earlier in the dim light condition (mean latency to shelter dim: 3.1 s, bright: 5.4 s; Figure 2H). On the other hand, *P. maniculatus* mice were more likely to enter the shelter under dim conditions (dim: 18/ 20 mice; bright: 6/11), but did so at similar latencies as during the bright light condition (Figure 2H). The characteristics of darts were different both between the species and under the different lighting conditions (Figure 2J-K). Under dim light conditions, darts to safety were more stimulus-locked, while also containing more intermediate sprints (Figure 2J; Suppl. Fig. 2E). *P. polionotus* mice produce d more stimulus-locked darts to safety than its sister-species, *P. maniculatus* (Figure 2K). In summary, while the specifics of the behaviors were different between the three species, each of them responded earlier to a black looming stimulus under dim compared to bright conditions, and displayed a more vigorous behavior characterized by switches to darts and increased escape speeds.

### Reduced levels of habituation under dim light conditions

Animals, including mice, in the absence of consequence or reward tend to habituate to repeated exposure to the same stimulus^19,20^. To test whether ambient light had an impact on habituation, we exposed separate cohorts of *P. polionotus* (N = 10) and *M. musculus* (N = 7) to sets of 10 looms at every entry of the threat zone until the animal did not show any responses or remained in the shelter or corner for more than 10 minutes. At bright light, we observed the previously reported habituation to repeated looms in *P. polionotus* where the percentage of tested and reacting animals decreased after a few repetitions, which was much less pronounced under dim light conditions (Figure 2L). Only for one *P. polionotus* the experiment at bright light was ended because it remained in the shelter for >10 min, all other individual stopped reacting to the stimulus over time. In *M. musculus* the percentage of responding mice across stimulus repetitions decreased as well at bright light, however, not because the mice ignored the stimulus, but further testing was not possible because they remained in the shelter for too long (Suppl. Fig. 2D). These prolonged, looming-induced shelter visits (>10 min) were further enhanced under dim light conditions where some individuals remained in the shelter already after the first presentation (Figure 2L). Although the precise effects of dim light differed between the two species, these results suggest that darker ambient light not only increases responsivity of mice to looming predators, but also decreases habituation to such stimuli.

### Perceived threat level of visual stimuli changes with lighting condition

It has been regularly reported that, in comparison to black expanding/looming discs, white expanding/looming and dimming discs tend to trigger behaviors consistent with a reduced risk level^5,21,22^. It appears that both the contrast and the expansion property are important to elicit aversive reactions and stimuli with out these properties are often ignored, and sometimes approached and examined^5^. Having observed stronger reactions to dark looming stimuli under dim light conditions, we wondered whether changing the ambient light condition would also impact behavioral responses to putatively “non-threatening” stimuli. We hence exposed mice to white looming and dimming stimuli under dim and bright light conditions (Figure 3A). In accordance with previous reports, we observed that, in bright conditions, all three species exhibited only occasional escape-like aversive behaviors in response to a white looming *(M. musculus:* 27% escape or darts; *P. maniculatus*: 8%; *P. polionotus*: 9%) or dimming stimulus *(M. musculus:* 20% escape or dart; *P. maniculatus*: 36%; *P. polionotus*: 28%, Figure 3B). However, when exposing mice to the same stimuli under dim conditions, both *P. polionotus* and *M. musculus* exhibited strong aversive behaviors (Figure 3B-C). White looming disks caused increases in escape or dart behavior in *M. musculus* (dim vs bright: 37% vs 20% of all trials) and in *P. polionotus* (31% vs 9%). Similarly, the presentation of a dimming disk under dim light conditions led to pronounced shifts from no reaction to aversive behaviors in *M. musculus* (escape: 50% dim vs 10% bright; see also Suppl. Figure 3A). In contrast, *P. maniculatus* does not follow this trend but appears to rather lose responses to dimming stimuli (10% vs 36% of escape/dart behavior under dim/bright light). In addition to stopping, escaping and darts, we also observed changes in ‘weak responses’ which include brief startles and rearing behavior, which were generally more prominent under brighter conditions. Interestingly, for overhead sweeping disks, considered a threat-like stimulus mimicking an overhead cruising predator, we found similar species-specific effects instead of a general light-dependent trend (Suppl. Fig. 3B). In conclusion, in contrast to general light-specific responses to looming stimuli, we find species-specific changes in aversive reactions to ‘non-threat’ stimuli that depend on the lighting conditions. These experiments further demonstrate that stimuli previously considered to be non-threatening controls reliably elicit aversive reactions in mice under dim ambient lighting conditions.

**Fig. 3:**
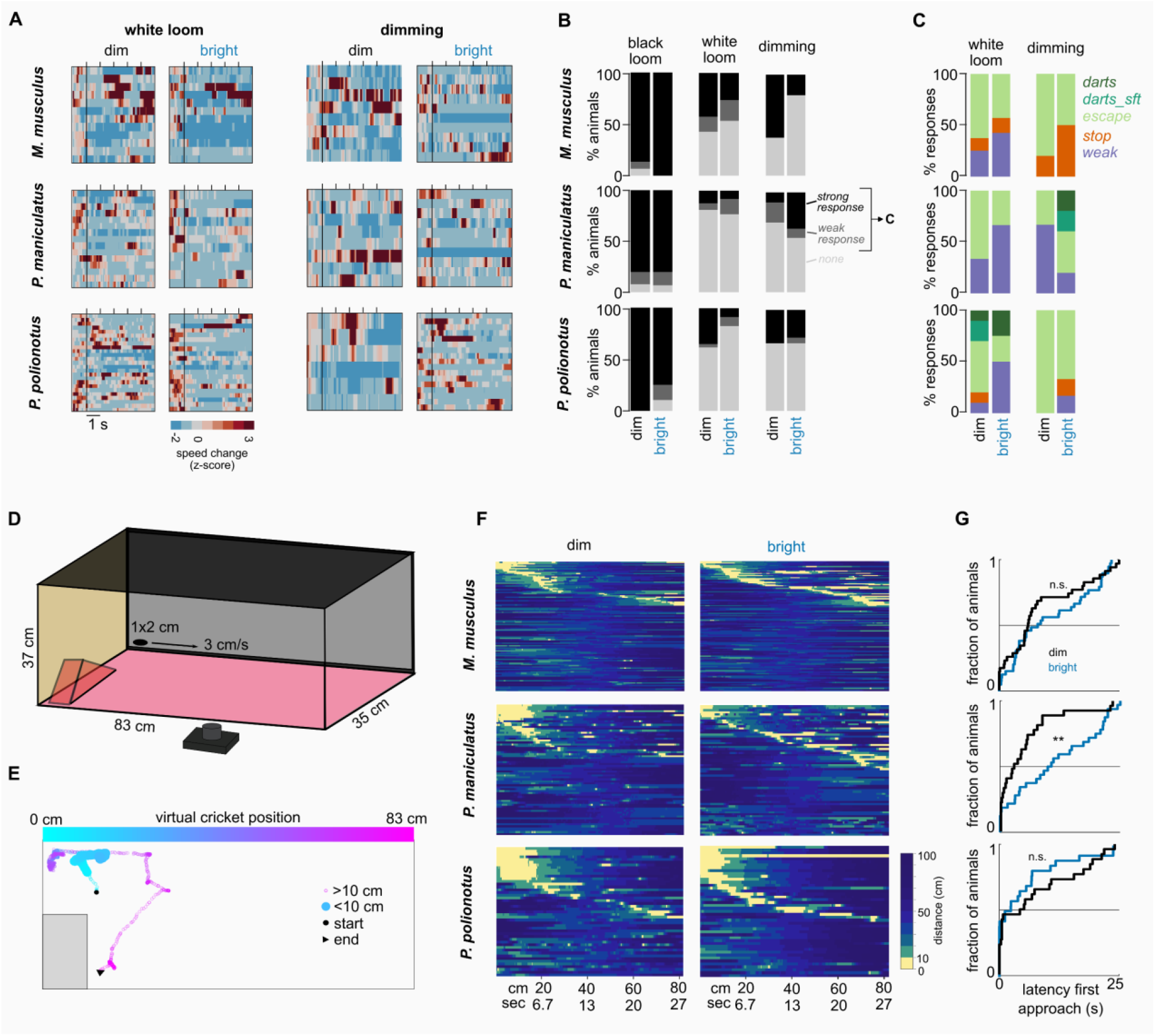
Species-specific responses to non-threat and virtual cricket stimuli. (**A**) Speed heatmaps during presentations of white looming and dimming stimuli. (**B**) Number of mice responding to black looming and non-threat stimuli (white looming and dimming). Strong responses include darts, escape and stopping. (**C**) Distribution of type of behaviors for weak and strong responses (black and dark gray in B). (**D**) Behavior setup with virtual cricket stimulus. (**E**) Example trajectory of a mouse reacting to the virtual cricket (Suppl. Video 9). Color indicates position and time. (**F**) Distance of the mouse to the cricket during all virtual cricket presentations for *Mus, P. maniculatus* and *P. polionotus* under dim and bright light conditions. Presentations are sorted by the time of the first approach. The cricket moved from the shelter side (0 cm) to the opposite short side (83 cm) at 3 cm/s. Number of mice for *Mus* (dim/bright): N = 12/12 (N = 11/12 with at least 1 approach, N = 7/11 with at least 2 approaches); *P. maniculatus*: N = 7/7 (N = 7/7, N = 7/7); *P. polionotus*: N = 7/7 (N = 7/7, N = 7/6). (G) Latency to the first approach. ** p < 0.01 Permutation Test.

### Approach behaviors in dim and bright environments

Hunting, like the avoidance behaviors discussed above, contains a number of innate components including the detection and approach phases^6^. To test whether these behaviors are affected by ambient light levels, we exposed mice to small ellipses (Figure 3D) that when moving slowly at ground level have been shown to trigger orienting and approach behaviors, similar to those displayed by mice hunting a cricket^6^. The moving ellipse was presented each time the animal was close to the side from where the ellipse appeared and faced the screen. Approaches were defined as movements resulting in bringing the center of body mass within 10 cm of the ellipse (Figure 3E). All three species showed clear approach behaviors in both dim and bright lighting conditions (Figure 3F, Suppl. Video 9-11). We observed a small trend of less approaches in the dim light condition (Suppl. Fig. 3C; mean successful presentations (dim vs. bright): *M. musculus*: 33% vs. 40%; *P. maniculatus*: 44% vs. 53%; *P. polionotus*: 56% vs. 62%). The duration of following and engagement with the moving ellipse was not affected by the change in ambient light, with *P. polionotus* consistently being the most persistent hunters (Suppl. Fig. 3D). However, the latency to initiating an approach differed between the two lighting conditions in a species-specific manner (Figure 3G). *P. polionotus* had a tendency to approach the moving ellipse earlier under bright conditions (median latency (dim/bright): 4.9 s / 1.9 s), whereas *P. maniculatus* approached the ellipse earlier under dim conditions (2.9 s / 10.7 s). In *M. musculus*, we observed a similar trend as in *P. maniculatus*, with shorter latencies under darker ambient light (6.3 s / 8.4 s). Overall, we found that ambient light had the strongest effect on *P. maniculatus* approach behavior with earlier approaches under dim light conditions, while it had little effect on the other two species.

### Light level affects pre-stimulus exploratory behavior

Finally, we asked whether spontaneous exploration in the arena differed between the two ambient light levels. To address this, we first compared the speed profiles of the mice during the 5 seconds immediately preceding each stimulus presentation. This corresponded with the time shortly before the mice entered the threat zone. We observed a general preference for movement at higher speeds during this pre-stimulus period of exploration in the dim condition for all three species (Figure 4A). Indeed, light level was the best predictor of pre-stimulus speed (generalized linear model: pre_stimulus_speed ∼ 1 + light_level + behavior + species; light_level p = 0.0003, behavior p = 0.18, species p = 0.28). Other characteristics describing the behavior before the stimulus, including time spent in the center, threat zone or shelter, were better explained by the species identity and not by the light level (generalized linear model for time spent in the center: species p < 0.0001, other p > 0.1; time spent in threat zone: species p < 0.0001, other p > 0.2; time spent in shelter: species p = 0.0008, light_level p = 0.2, behavior p = 0.03). In a subset of *P. polionotus* mice we also recorded their foraging behavior between stimulus presentations (N=10 mice, 2.5 min recordings before each stimulus). Interestingly, the majority of these mice moved slower under dim light levels (Figure 4B), even though this subset of mice showed the same trend of higher pre-stimulus speed under dim conditions as the other individuals (Figure 4C). Similar results were obtained with a separate cohort of *M. musculus* recorded at a different institute (Suppl. Figure 4). Taken together, this suggests that while mice forage more slowly or less in dimly lit environments, when they explore spaces associated with an increased level of risk they do so at higher speeds.

**Fig. 4:**
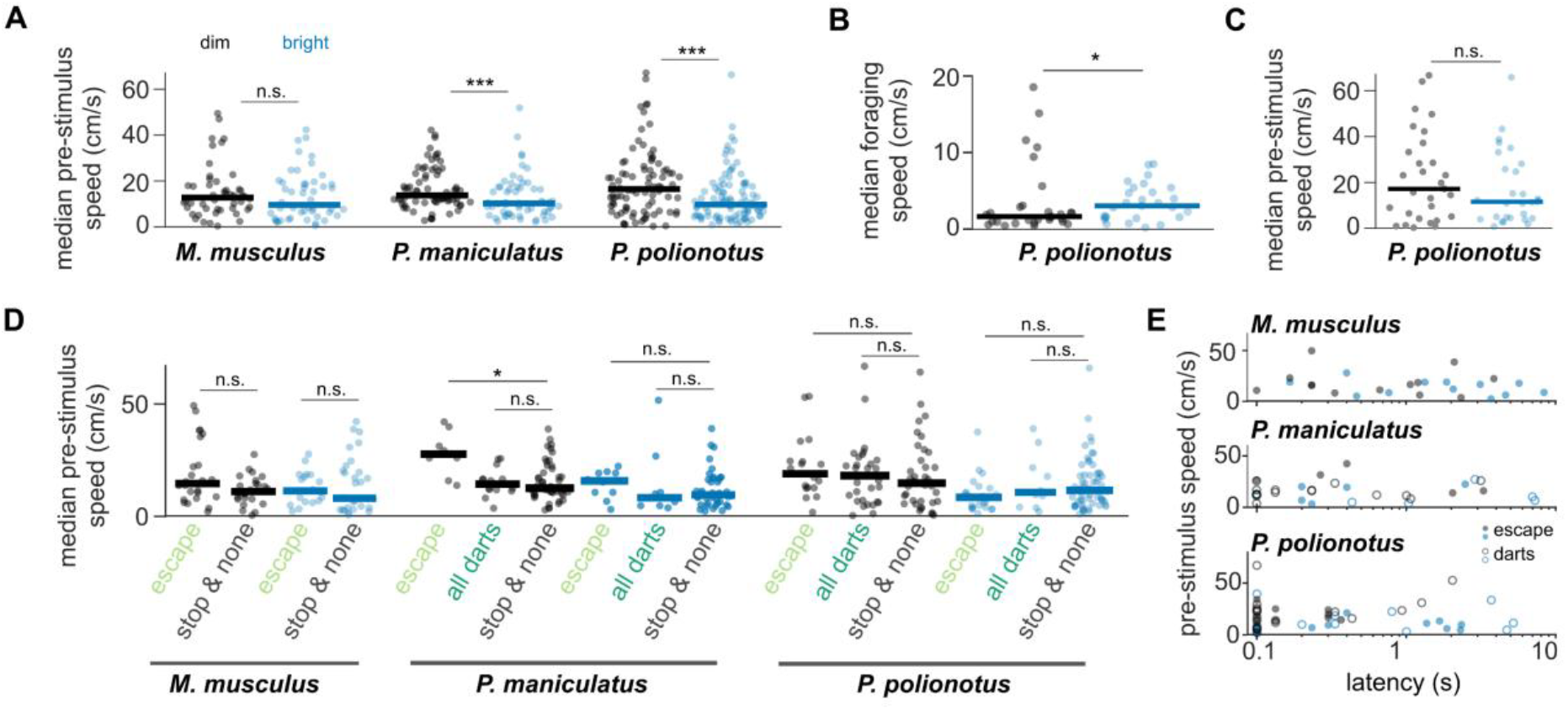
Ambient light predicts pre-stimulus behavior. (**A**) Median speed during the 5 seconds right before each stimulus onset (pre-stimulus speed) for each species and both light levels. Lines indicate medians. (**B**) Median speed during 2.5 minutes between each stimulus presentation for a subset of *P. polionotus* mice (N = 10). (**C**) Pre-stimulus speed for the same mice as in B. (**D**) Pre-stimulus speed for black looming stimuli only, separated by subsequent elicited behavior. (**E**) Pre-stimulus speed and latency to running onset for escape (filled) and dart (open) trials at dim (black) and bright (blue) light levels. * p < 0.05, ** p < 0.01 *** p < 0.001 Wilcoxon Ranksum Test corrected for multiple comparisons.

### Light level but not pre-stimulus activity predicts behavior

We wondered whether the higher speeds observed during the pre-stimulus phase in the dim condition might facilitate the initiation of sprints during visually triggered escapes and darts. We found no consistent relationship between the running speed of mice during the pre-stimulus phase of the experiment and the behaviors triggered by the visual stimulus; except for *P. maniculatus* mice that showed a trend for running faster before an escape but not before darts (Figure 4D). The weak relationship between pre-stimulus running and behavior was confirmed using a linear model where the type of elicited behavior is best predicted by ambient light levels (pre_stimulus_speed p = 0.18, light_level p = 0.02, species p = 0.06). Since the main change between looming responses at different light levels was the onset latency (Figure 2), we tested whether pre-stimulus speed would predict escape latency rather than behavior type. In all three species, median pre-stimulus speed did not correlate with escape latency of the subsequent looming stimulus (Figure 4E; *M. musculus*: r = -0.23, p = 0.3, *P. maniculatus*: r = -0.1, p = 0.6, *P. polionotus*: r = -0.03, p = 0.8). Accordingly, a linear model showed that light level is the dominant predictor of escape latency, followed by species, but confirmed that there is no relationship between pre-stimulus speed and escape latency (pre_stimulus_speed p = 0.54, light_level p = 0.00005, species p = 0.004). In conclusion, overall pre-stimulus running speed as well as stimulus-triggered behavior type and latency are strongly dependent on the ambient light levels in all three species. However, pre-stimulus running speed before the loom ing stimulus is not predictive of the likelihood or latency of escape (see also Figure 2).

## DISCUSSION

Here we demonstrate that ambient light levels shape the expression of innate avoidance and approach behaviors in three different species of mice. We identified both conserved and species-specific behavioral adaptations to visual threats and opportunities, revealing that lighting conditions can dynamically modulate inherited behavioral programs. In particular, all three species responded earlier and more vigorously to overhead predator-like stimuli under moonlight-like conditions, while species differed in their responses to non-threatening or prey-like cues. Importantly, we show that pre-stimulus exploratory behaviors are influenced by ambient light but do not predict subsequent visually evoked actions, suggesting that changes in behavior arise from central modulation of sensory-motor pathways rather than differences in arousal or activity. Together, these findings reveal that environmental light conditions are a critical, yet often overlooked, factor shaping innate behaviors, and highlight the need for careful consideration of naturalistic environmental variables when interpreting behavioral and neural data. More broadly, our work provides a tractable framework for studying how inherited neural circuits integrate dynamic environmental contexts to flexibly modulate survival behaviors.

In addition to carnivorous mammals, birds, including night-active owls, are common predators of both Eurasian and American mice^8–10,23^. Owls have been found to be more efficient hunters during a full-moon9, which corresponds to our dim or moonlight condition. The efficiency of their predators under these lighting conditions might be the reason for the fast onset and more vigorous reaction of all three rodent species to overhead black looming stimuli. In *M. musculus*, we observed shelter-directed escapes as the main behavior to an approaching threat. While this is the shortest way to safety it is also predictable. Many species from insects to mammals choose instead more varied and erratic (‘protean’) escape strategies^24^ that result in them being more difficult to catch^25,26^. Moths, for instance, show directed escape when exposed to distant ultrasonic stimuli, but switch to chaotic flight paths when threats are imminent^27^. We observed a similar strategy in the two *Peromyscus* species, which displayed darting behavior throughout the setup, in particular at low light levels. Especially in *P. polionotus*, whose natural habitat are exposed sandy dunes, erratic runs may be the best strategy to avoid predation.

We find that moonlight leads mice to be more cautious, particularly *M. musculus* and *P. polionotus*. This is reflected in a lower threshold to trigger avoidance responses to “non-threatening” stimuli, slower and less expansive exploration, and less habituation to repeated presentation of looming stimuli at these light levels. This apparent higher level of awareness of danger in dim conditions might also be why virtual crickets were approached slightly less often and for shorter periods of time than during daylight conditions. Similar to our results in *P. polionotus*, spontaneous foraging has been found to decrease during moonlit nights in wild populations of *Peromyscus leucopus*^13^ and *Peromyscus maniculatus*^9,28^ (but see^29^). However, it is important to note that all these studies compare foraging at moonlight to even darker conditions and not to daylight. While in the past mice may not have been very active under the low photopic light levels tested in our experiments as well as in most other laboratory studies, today’s light pollution in cities increases the likelihood of mice encountering brightly lit environments even during the night.

The neural mechanisms that allow modulation of fast and reflex-like innate behaviors are largely unknown. Innate aversive and orienting behaviors are mediated by circuits through the superior colliculus, which receives behavior-driving inputs from the retina, but also dozens of inputs from across the brain^30,31^. Several potential mechanisms could modulate those circuits depending on ambient light. For instance, ambient light affects visual encoding beginning at the level of retinal output neurons^32–34^. Furthermore, direct or indirect melanopsin signaling (the retinal light integrator) to the superior colliculus may affect visuo-motor encoding^35^. Recent publications have shown that decreased firing of colliculus-projecting GABAergic neurons in the ventrolateral geniculate nucleus (vLGN) increases aversive reactions and that vLGN activity increases with ambient light^36,37^. This poses one possible mechanism for the systematic behavior changes that we describe here, where we might see stronger escape at dim light levels due to decreased vLGN inhibition to the colliculus. However, how these circuits act in different species, especially in the light of the species-specific changes of responses to “non-threatening” and appetitive stimuli, remains unknown.

Animals will experience different ambient light levels both because of changes in the time of day, but also because of different weather conditions, changing seasons, when entering or leaving their burrow or when switching between open and shaded areas.

While circadian rhythm affects a variety of rodent behaviors (forced-swim test performance, cutaneous sensitivity, long-term memory, vigilance, exploration in elevated mazes, conditioned contextual fear behavior and spontaneous escaping of small enclosures)^38–42^, some behaviors, including open field exploration or active avoidance of electric shocks do not appear to change with the time of day^38^. To our knowledge, the effect of circadian rhythm on innate aversive or predatory behaviors has not been studied, and it remains to be determined whether the light-dependent behaviors we describe here are due to related circadian time points or rather due to other associations with these light levels.

In conclusion, we find that innate behaviors are not rigid but can be adaptively modulated by ambient light conditions, and that these changes are inherited rather than learned. This insight has major implications for interpreting behavioral and neural responses to visual stimuli in laboratory settings, where lighting conditions often differ substantially from natural environments. Furthermore, these results show that evolutionary pressures have shaped both general and species-specific strategies for balancing foraging and predator avoidance under different illumination levels. Given the central role of subcortical circuits in driving innate behaviors, this work provides a novel, pragmatic framework for dissecting the neural circuit mechanisms that enable flexible modulation of hard-wired, survival-critical behaviors by environmental cues. More generally, our findings underscore the importance of integrating real-world conditions into experimental design to better understand how innate behaviors evolve and operate in the natural world.

## ACKNOWLEDGEMENTS

We would like to thank Hopi Hoekstra (Harvard University) for providing us with *Peromyscus* breeding pairs and for advice with *Peromyscus* handling and experimentation, and Jennifer Hoy (University of Reno) for advice with the cricket stimulus and valuable feedback. Further, we thank Anna Chrzanowska and Bram Nuttin (both NERF) for helping with the behavior setup and with DeepLabCut, Marco Gigante for building the setup at SISSA, and the Farrow Lab and Bonin Lab (both NERF) as well as Eugenio Piasini (SISSA) for helpful comments. This project has received funding from the European Union’s Horizon 2020 research and innovation programme under the Marie Skłodowska-Curie grant agreement No 665501 and from the European Research Council (ERC) (Grant agreement No. 101075848) as well as from FWO (12S7917N and 12S7920N) to KR. The views and opinions expressed are solely those of the authors and do not necessarily reflect those of the European Union, nor can the European Union be held responsible for them. KF is supported by the FWO (G094616N and G091719N) and the NIH (1R01EY032101).

## AUTHOR CONTRIBUTIONS

Conceiving the study: KR, KF, SDB

Pilot experiments: SDB, LT

Data collection & processing: KR, RV, BC, PYL, LT

Analysis: KR, KF

Manuscript writing: KR, KF

## MATERIAL & METHODS

### Key resources table

| REAGENT or RESOURCE | SOURCE | IDENTIFIER |
| --- | --- | --- |
| **Experimental models: Organisms/strains** |  |  |
| *Mus musculus*: C57BL/6 | The Jackson Laboratory | JAX: 00664 |
| *Peromyscus maniculatus bairdii* | Hopi Hoekstra, Harvard University | BW |
| *Peromyscus polionotus subgriseus* | Hopi Hoekstra, Harvard University | PO |
| **Software and algorithms** |  |  |
| DeepLabCut | Mathis et al., 2018; Nath et al., 2019 | http://www.mackenziemathislab.org/deeplabcut |
| Python Jupyter Notebook | Python Software Foundation | https://jupyter.org/<br>https://www.python.org/ |
| Matlab | The MathWorks, Inc. | https://www.mathworks.com/ |
| Dabest | Ho et al. 2019 | https://acclab.github.io/DABEST-python/ |
| Psychtoolbox | Brainard 1997 |  |
| Pylon | Basler | https://www.baslerweb.com/en/software/pylon |
| Bonsai | Lopes & Monteiro, 2021 | https://bonsai-rx.org/ |
| BonVision | Lopes et al. 2021 | https://bonvision.github.io/ |
| InkScape |  | https://inkscape.org/ |
| ImageJ | Schneider et al., 2012 | https://imagej.net/software/fiji/ |
| **Other** |  |  |
| Cameras | Basler | acA1300-60g mNIR |
| Objective bottom camera | Ricoh Lens | FL-CC0614A-2M |
| Objective top camera | Computar | 8mm 1:1.4 2/3 |
| LED light | Solarox | LED 850nm |
| Display monitors | LG | 34UM69G-B |
| Neutral density filters | Lee | 211 0.9 ND |

### Resource availability

#### Materials availability

This study did not generate new unique reagents.

#### Data and code availability

X- and y-coordinates for each animal, stimulus and condition will be made available on a public repository upon publication.

Code used to create the figures will be made available on a public repository upon publication.

### Experimental Details

#### Animals

For the main data sets, we used 26 *Mus musculus* (C57BL/6J, Jax strain 00664), 41 + 10 *Peromyscus polionotus subgriseus* and 39 *Peromyscus maniculatus bairdii* (both obtained from Hopi Hoekstra, Harvard University) of both sexes at the age of at least 2 months to up to 6 months (*Mus*) or 9 months (*Peromyscus*). Mice were housed in the facilities of Neuroelectronics Research Flanders under a 12/12 dark/light cycle at 23°C and with access to standard mouse food and water ad libitum. All mice were housed in groups of at least 2 and maximal 5 mice. Mice were only briefly handled during the biweekly cage changes, but not during experiments. An additional 7 C57BL/6J mice and 15 Gad2-Cre^+/+^ (Jax strain JAX10802) mice housed under comparable conditions in the facilities at Scuola Internazionale Superiore di Studi Avanzati were used for the data presented in Figure 1L and Suppl. Figure 2D (C57BL/6J) and Suppl. Figure 2A, 3A and 4 (Gad2-Cre^+/+^). All experiments were approved by the Animal Ethics Committee of KU Leuven (protocol number 165/2018 and 219/2017) or the Italian Ministry of Health (authorization number 703/2023-PR).

#### Behavior setup

The experimental setup consisted of a box (35 x 82.5 x 37 cm) with a red Plexiglas floor and a computer monitor (LG34UM69G) on the top and on one long side. The shelter, consisting of a red Plexiglas triangle, was placed in one corner. The entrance box to the setup was in the opposite corner of the same short side. One camera was placed under the setup to film the animal’s behavior at 30-67 Hz under the control of pylon software (Basler) or the BonVision^43^ package of Bonsai^44^. A second camera was placed on the top of the shelter-containing short side to detect the onset of the cricket stimulus. Deep red lights (850 nm) were placed under and at the top edge of the setup.

#### Visual stimulation

Visual stimuli were generated using Octave (GNU Octave) and Psychtoolbox^45,46^ or BonVision, and presented on either the top or side screen. Stimuli consisted of (1) a black looming disk shown on a grey background and expanding from 2 to 20° in 500 ms, followed by 500 ms of grey screen. The stimulus was shown in the center of the top screen and repeated 10x for each session. (2) a sweeping black disk of 2° visual angle moving at 21°/s in the center of the top screen along the long side from the shelter-opposite side towards the shelter. (3) a white looming disk with the same specifications as the black looming disk, but shown on a black background. The stimulus was repeated 5x. (4) a dimming disk of a size of 20° which dimmed from grey to black within 500 ms, followed by 500 ms of grey screen. The stimulus was repeated 5x. (5) a small ellipse of the size of 1 x 2 cm moving at 3 cm/s along the bottom of the side screen from the shelter towards to opposite side. The screen brightness (grey) was 128 lux, corresponding to a dim photopic light level. To achieve dim light conditions, three neutral density foils were taped to the screen to decrease the brightness to ∼0.2 lux (mesopic).

#### Experimental procedure

Mice were moved from the cage into the setup using a metal grid or transport box which they entered and left on their own. Mice were not touched by the experimenter before or during the experimental sessions. After 10 minutes (20 minutes for dim conditions) of acclimatization to the setup, the first visual stimulus was triggered as soon as the animal entered the farthest 1/3 of the setup opposite of the shelter (threat zone’). The order of stimuli was randomly chosen and each overhead stimulus was shown maximally once per session. Between each stimulus presentation, we waited at least 30-60 seconds (exact duration was randomly chosen). After this time, the next stimulus was triggered at the next entry into the threat zone. Stimuli were either triggered manually (using Matlab and Pylon) or automatically using online detection of threat zone entry (Bonsai). If the animal remained in the shelter or a corner for a prolonged time, stimuli that could not be presented yet were presented in a second session on another day. Approximately half of the mice were first exposed to moonlight conditions and the other half to daylight conditions. There were at least 5 days between the two different conditions. Cricket stimuli were shown in separate sessions from the threat stimuli and were repeated whenever the animal was facing the screen until the animal showed at least two approaches. Experiments were performed between 8 am and 5 pm (light on in the animal facility: 7 am to 7 pm).

#### Data processing

DeepLabCut^15,16^ was trained on the acquired videos to extract different positions on the mice including center of the body, paws, head, nose and tail (resnet50, 100000 iterations). Consensus among the different body parts was used to calculate the center of mass of the animal. Similarly, the stimulus position was tracked throughout the video. We used in-house written Python code to display and correct the extraction of the animal position 10 s before and during the stimulus. From the x and y position of the mice’ center of mass we calculated the position of the animal at each frame as sqrt(x^2 + y^2) and the speed by taking the difference of those positions. The speed data was then resampled to 30 Hz, smoothed with a smoothing window of 5 data points, and transformed into cm/s based on the pixel resolution and the setup size.

#### Parameter extraction

In-house written Matlab code was used to extract the following parameters from the speed and position data and to produce graphs. Figures, including setup schematics, were assembled in InkScape. DABEST statistics were performed in Python.

##### Percent speed change

To visualize changes in the behavior after stimulus onset across species with very different baseline and evoked running speed, we calculated a z-score of the speed. The z-score was based on the mean and standard deviation of the speed in the 10 s before stimulus onset.

##### Distance to shelter and shelter entry

The distance to shelter was calculated for each recorded frame by subtracting the edge x- and y-coordinates of the shelter from the x- and y-coordinates of the animal. To account for small movements of the shelter and the non-rectangular shape, shelter entry was defined as a distance of <5 cm from the shelter corners.

##### Running bouts

Running was defined as speeds > 30 cm/s for at least 3 consecutive frames. This threshold was determined as the 97.5-percentile of baseline running speeds in *Mus* (see Suppl. Figure 1D). In Peromyscus, this threshold corresponds to the 92-percentile.

##### Latency to running

Latency was defined as the time between stimulus onset and onset of a running bout.

##### Number, latency and duration of cricket approach

Cricket approach was defined as distance of <10 cm between the animal and the cricket stimulus for at least 10 frames (0.3 seconds).

##### Maximal running speed

For validation of manually scored trials (Figure 1K), maximal running speed was defined as the maximal speed in the first 5 seconds after stimulus onset.

##### Stopping time

For validation of manually scored trials (Figure 1K), the maximal number of consecutive frames below 5 cm/s was taken as the stopping time.

##### Power spectrum of Fourier transform

To compute the Fourier transform, speed traces during the stimulus presentation were first detrended (the best straight-fit line was subtracted from the data) using the matlab function ‘detrend’. Then, the Fast Fourier Transform Y was calculated using the function ‘fft’. The Power spectrum was subsequently computed using the following steps:

P2 = abs(Y/L) with L being the length of the speed traces to create a two-sided spectrum

P1 = P2(:,1:L/2+1) to create a single-sided spectrum P1(:,2:end-1) = 2*P1(:,2:end-1) to calculate the root mean square amplitude per frequency

P1 = P1.^2 to calculate the mean squared amplitude

##### Pre-stimulus speed

Pre-stimulus speed was defined as the median speed during the 5 seconds preceding a stimulus presentation.

##### Foraging speed

Foraging speed was defined as the median speed during the 2.5 minutes preceding a visual stimulus.

##### Percent time in danger zone, center, shelter

For pre-stimulus behavior analysis, the danger zone was defined as the half of the setup on the opposite side of the shelter; the center was defined as the central half of the setup (X: length/4:length/(3/4); Y: width/4:width/(3/4)); and shelter zone was defined as a rectangle of 23 cm side length starting at the shelter corner of the setup. For each parameter, the time spent in these parts of the setup was calculated as a percentage of the pre-stimulus (5 sec) or foraging time (2.5 min).

##### Safe zones

To identify darts that could have been included in previous definitions of escape behavior (speed threshold plus ending in shelter), we determined whether dart trials ended in a safe zone. Safe zones were defined as the zones along the short ends with a 20cm width on the shelter side (i.e., far from the stimulus eliciting zone) and with a 10cm width on the far end (see Suppl. Figure 1B).

#### Manual scoring

Videos around each stimulus presentation were watched and scored manually into the main reaction to the stimulus. The 5 behavior categories were defined as follows:

##### Stopping

Complete halt or freezing (complete immobility of all body parts).

##### Escaping

One strong, immediate run towards the shelter or in another direction followed by stopping. Trials where the mice stopped very briefly and then immediately ran away from the stimulus were also counted as escapes.

##### Darts

Multiple instances of stopping, sprinting, stopping into different directions for each sprint. Darts were further split into those that ended in safety (within designated safe zones during the last second of the looming presentations) and darts that did not end in safety.

##### Weak responses

Very brief startles, not followed by runs; and approaching movements including rearing towards the stimulus.

##### None

The stimulus was in the visual field of the animal, but none of the reactions described above could be observed.

#### Exclusion criteria

We excluded all trials where the stimulus was not in the animal’s visual field. This could happen because the experimenter triggered the stimulus too late or because the animal had left the threat zone very quickly after entering it and before the stimulus appeared. Some additional trials had to be excluded due to technical issues that were only detected after the experiment such as incorrect lightning or stimulus parameters. Finally, some individuals did not complete all stimuli due to prolonged hiding in the shelter during multiple sessions.

#### Statistical analysis

Statistical tests included Permutation Tests (performed using the ‘dabest’ package (https://acclab.github.io/DABEST-python/)^47^, Wilcoxon Ranksum tests (‘ranksum’ function in Matlab) with Bonferroni correction for multiple comparisons where indicated, and cross-correlation calculations:

Average speed plots (Figure 2): xcorr’ in Matlab was used to calculate cross-correlations between the median speed during the looming stimulus presentation and the stimulus. The stimulus was represented as a sinusoidal function with the 1 Hz frequency of the looming stimulus.

Dart behavior (Figure 1 and Figure 2): A matrix was created with 0s and 1s where 1 indicates the presence of a running bout. The correlation between this matrix and a stimulus vector with 0s and 1s where 1 indicates the onset of each stimulus was calculated using the ‘xcorr’ function.

Escape latency vs. pre-stimulus speed (Figure 4): Correlation coefficients and corresponding p-values between escape latency and pre-stimulus speed were calculated using the ‘corrcoef’ function.

