## Supplemental Figures for "Ambient Light Impacts Innate Behaviors of New-World and Old-World Mice"

### Supplementary Figures

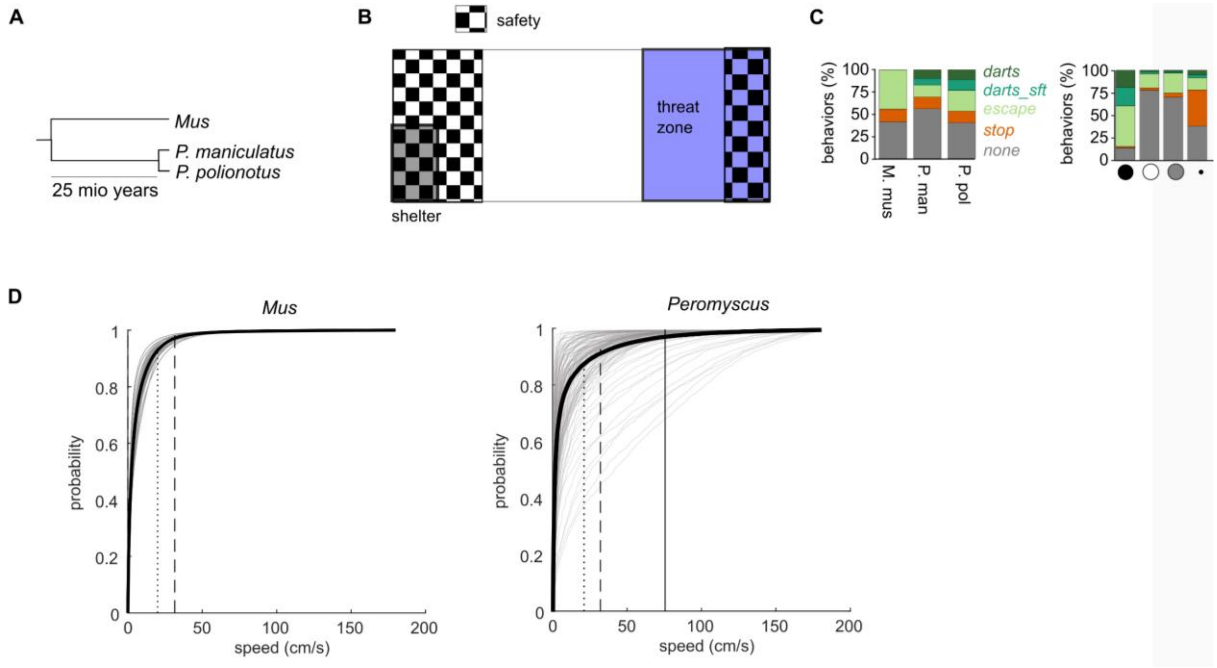


**Suppl. Fig. 1: Behavior sequences for manually categorized behavior classes.** (A) Phylogenetic relationship of the three tested rodent species. (B) Indication of safe zones in the setup relative to shelter and threat zone. (C) Percent behaviors across all stimuli and light levels (left); and percent behaviors across all species and light levels (right). (D) Cumulative distribution of movement speeds during 2.5-minute spontaneous foraging phases for a subset of *Mus* and *P. polionotus* individuals. Gray lines are individual trials, black line depicts the median. Corse dashed lines indicate 97.5% quantile of median *Mus* speed (30 cm/s). Fine dashed line indicates 20 cm/s. Solid vertical line indicates 97.5% quantile of median *P. polionotus* speed (74 cm/s). Speed distributions were less consistent across *Peromyscus* individuals, and the *Mus* 97.5% cut-off was used as an escape threshold for further analysis (analysis adapted from Lenzi et al. 2022).

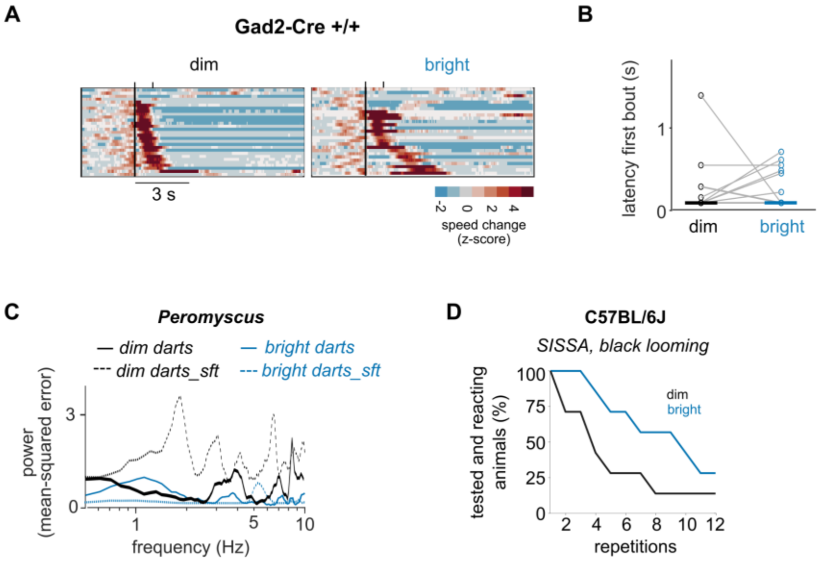


**Suppl. Fig. 2: Additional analysis of black looming responses.** (**A**) Looming responses of 15 Gad2-Cre$^{+/+}$ animals at dim (29 trials) and bright light levels (27 trials). Sorted by running latency. Experiments were performed at a different institute than for Figure 2. (**B**) Latency changes of onset of first running bout for the faster reaction per animal from A. (**C**) Power spectrum of Fourier transform of darts in response to the black looming stimulus for dim and bright light conditions separately. (**D**) Percentage of tested and responding mice (N=7 C57BL/6J tested at SISSA) for repeated sets of 10 looming stimuli. If mice could be tested (i.e., did not stay in the shelter for >10 min) they always escaped in response to the looming stimuli. The decrease in percentage was solely due to prolonged shelter visits, which were enhanced at dim light.

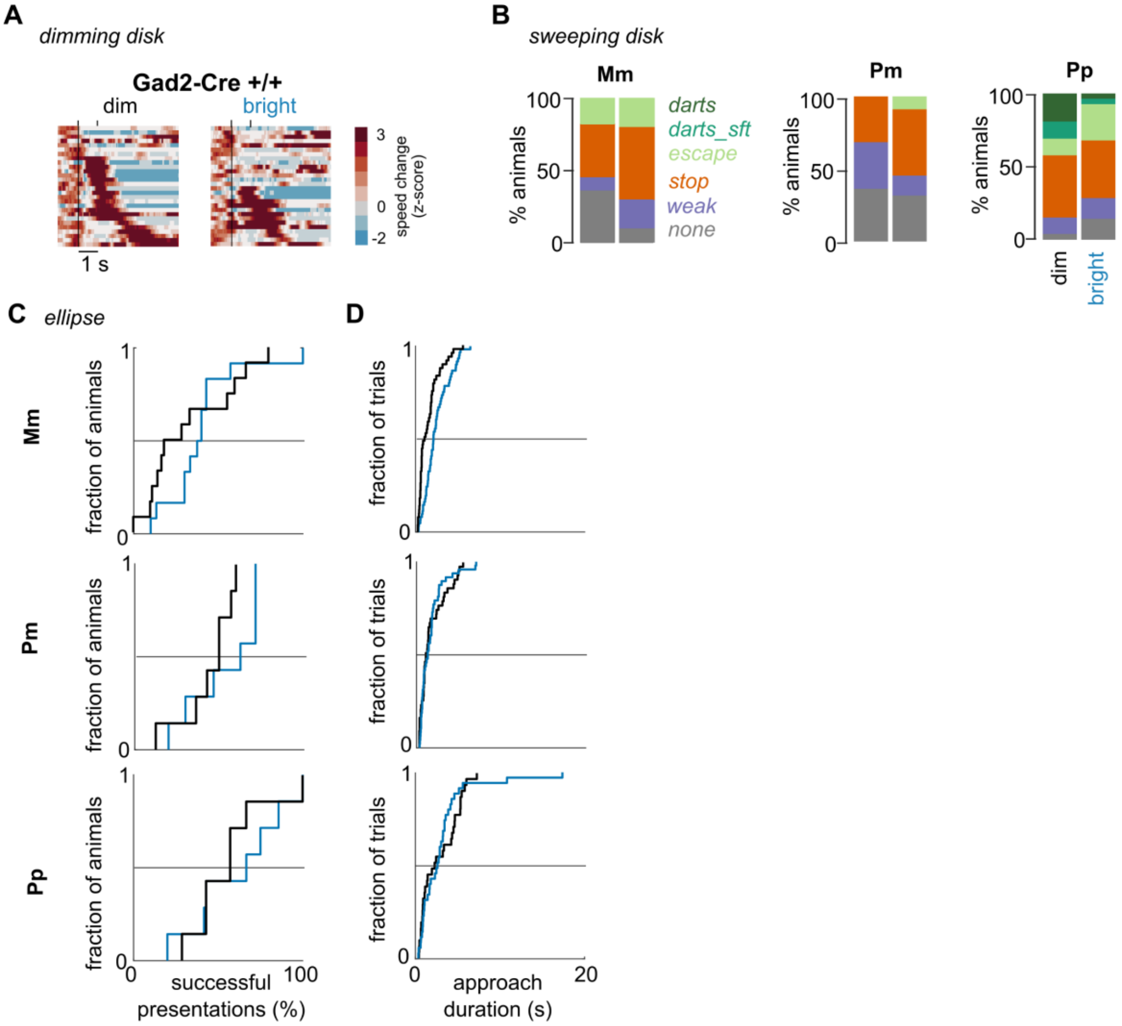


**Suppl. Fig. 3: Additional information about non-threat stimuli responses.** (**A**) *Mus musculus* responses to a dimming disk recorded at a different institute and in a different strain. This cohort tested in this different environment and setup displayed more responses to a dimming disk already at bright light than the cohort in Figure 3A, however, the same increase in responses under dim light can be observed (28 trials each from 14 animals; same cohort as in Suppl. Fig. 2A). (**B**) Behavior types in response to a small overhead sweeping disk mimicking an overhead cruising predator. (**C**) Cumulative distributions of the number of cricket-like stimulus presentations that led to an approach behavior. (**D**) Cumulative distributions of approach durations of the cricket-like stimulus.

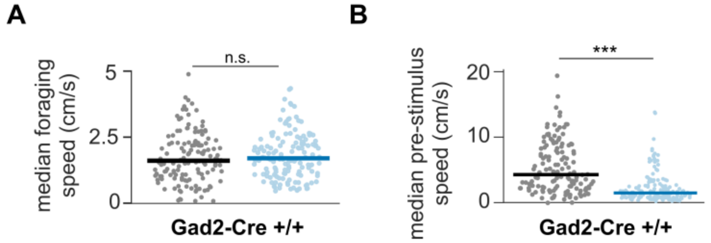


**Suppl. Fig. 4: Foraging and pre-stimulus speed in *M. musculus*.** (**A**) Foraging speed, i.e. locomotion speed in the 1.5 min between stimuli of any type of 14 separately tested *M. musculus* (same animals as in Suppl. Fig. 2A and 3A). Experiments were performed at SISSA. (B) Median pre-stimulus speed (5 s before stimulus onset) of the same animals as in A. Overall, the same trend as in *P. polionotus* (Figure 4B-C) can be observed with slightly higher foraging speeds but lower pre-stimulus speeds at bright light. *** $p < 0.001$ Wilcoxon Ranksum Test.
